# Metagenomic Immunoglobulin Sequencing (MIG-Seq) Exposes Patterns of IgA Antibody Binding in the Healthy Human Gut Microbiome

**DOI:** 10.1101/2023.11.21.568153

**Authors:** Matthew R. Olm, Sean P. Spencer, Evelyn Lemus Silva, Justin L. Sonnenburg

**Affiliations:** Department of Microbiology and Immunology, Stanford University School of Medicine, Stanford, CA, USA; Chan Zuckerberg Biohub, San Francisco, CA, USA; Center for Human Microbiome Studies, Stanford University School of Medicine, Stanford, CA, USA; Division of Gastroenterology and Hepatology, Stanford School of Medicine, Stanford, CA, 94305, USA

## Abstract

IgA, the most highly produced human antibody, is continually secreted into the gut to shape the intestinal microbiota. Methodological limitations have critically hindered defining which microbial strains are targeted by IgA and why. Here, we develop a new technique, Metagenomic Immunoglobulin Sequencing (MIG-Seq), and use it to determine IgA coating levels for thousands of gut microbiome strains in healthy humans. We find that microbes associated with both health and disease have higher levels of coating, and that microbial genes are highly predictive of IgA binding levels, with mucus degradation genes especially correlated with high binding. We find a significant reduction in replication rates among microbes bound by IgA, and demonstrate that IgA binding is more correlated with host immune status than traditional microbial abundance measures. This study introduces a powerful technique for assessing strain-level IgA binding in human stool, paving the way for deeper understanding of IgA-based host microbe interactions.

## Introduction

Immunoglobulin A (IgA) is the primary human antibody secreted from mucosal surfaces. More IgA is produced each day than all other antibody isotypes combined, and most of it is secreted into the intestinal lumen ^1,2^. Of all antibody secreting plasma cells present in the human body, an estimated 60% reside in the intestine to maintain high levels of IgA throughout the GI tract ^3^. Mounting evidence indicates that IgA plays a critical role in maintaining gut microbiome homeostasis. Patients with IgA deficiency have increased incidences of autoimmune disorders, infectious disease, and levels of gut bacteria translocating the intestinal barrier ^4–6^, and conditions including ulcerative colitis and Crohn’s disease are associated with distinct IgA binding capacities and specificities ^7–9^. A greater understanding of IgA dynamics in both health and disease states will facilitate the development of new therapeutics, including IgA-based therapies ^10^ for precision modulation of the gut microbiota.

Approximately one in eight bacteria (∼12-15%) in human stool are bound by IgA ^7^. The taxa most highly bound by IgA are Proteobacteria (a phylum enriched in opportunistic pathogens) ^1^ and bacteria associated with immunomodulation (such as *Bifidobacterium* and *Akkermansia*) ^4^. The specific microbial cell-surface antigens targeted by IgA via the adaptive immune system have been mapped in highly-controlled mouse studies ^11^, but it remains unclear why certain taxa are targeted over others in the human intestine. IgA binding can cause a wide range of consequences for microbes in the gut, most of which are inhibitory ^11–16^. However, IgA binding can increase the colonization capability of some intestinal bacteria (including *Bacteroides fragilis*) ^17^. McLoughlin *et al.* describe how, by anchoring microbes to mucus, IgA binding could increase or decrease microbial fitness by varying the mucus flow rate ^18^. The “enchained growth” model describes how polyreactive IgA could aggregate and clear fast-replicating bacterial pathogens while not affecting more slower-growing commensals ^19^. Evaluating these hypotheses requires simultaneous measurement of bacterial IgA binding levels, *in vivo* replication rates, and microbial genome content, which is not possible using presently available methods.

IgA-SEQ^TM^ ^7^, the current state-of-the-art technique to study IgA binding, uses a combination of bacterial fluorescence-activated cell sorting and 16S gene amplicon sequencing to deliver a survey of the bacteria bound or unbound by IgA *in vivo* ^20^. However, the method has several notable limitations that greatly hinder understanding of IgA biology due to its reliance on bacterial 16S rRNA ribosomal subunit amplicon sequencing (16S sequencing). First, 16S sequencing has approximately genus-level taxonomic resolution. Closely related bacterial strains often have different levels of IgA coating ^7^, but the resulting broad taxonomic assignments prevent all investigation of strain- and species-level analysis of IgA targeting. Second, 16S sequencing cannot directly measure the genomic content of bound organisms, only their taxonomic identity. This limitation has substantially hindered our understanding of the specific microbial genes and functions associated with IgA targeting. Third, while escape from IgA binding via genomic mutation is hypothesized to be at the heart of some human infectious diseases ^21^, the lack of full-genome sequence data precludes the SNP-level analyses needed to evaluate such hypotheses. Finally, 16S sequencing is incompatible with modern algorithms that estimate *in vivo* bacteria replication rates from high-throughput sequencing data ^22,23^. This incompatibility has prevented evaluation of the “enchained growth” hypothesis and how IgA binding affects bacterial replication rates. The development of a method that uses metagenomic shotgun sequencing to measure IgA binding, as opposed to 16S sequencing, would overcome these shortcomings and facilitate a detailed understanding of IgA dynamics in the human gut and other mucosal environments.

Here we present Metagenomic Immunoglobulin Sequencing (MIG-Seq), a technique that leverages metagenomic shotgun sequencing to characterize immunoglobulin binding at the strain level in human stool samples. While the MIG-Seq technique is theoretically applicable to all antibody isotypes, in this manuscript we focus on IgA binding (IgA MIG-Seq). Using this technique we develop a framework to classify strains into discrete categories of IgA binding, identify specific microbial genes associated with IgA binding, and test the fitness impacts of IgA binding. Further, using high resolution, multi-omic immune profiling data associated with the human stool samples, we discover that strain-level IgA binding levels are significantly more correlated with host immune status than typically measured abundance-based microbiome metrics. These findings emphasize the value of IgA MIG-Seq and highlight its potential to enable greater understanding of the host / microbe immune interface.

## Results

### Metagenomic Immunoglobulin Sequencing (MIG-Seq) method development

Several modifications were made to the IgA-SEQ^TM^ protocol ^7^ to adapt it to utilize shotgun metagenomic sequencing, most of which were aimed at increasing IgA-bound microbial cell biomass (**Figure 1A**). Instead of Fluorescence Activated Cell Sorting (FACS) we utilized Magnetic Cell Sorting (MCS), a higher-throughput technique that achieved roughly 60% purity in the IgA+ fraction (mean=58.3% ± 2.2%; n=33) (**Figure 1B, C) (Supplemental Figure S1)**. By conducting comparative analysis of both the native and IgA+ fractions, accurate measures of IgA binding can be calculated with this level of purity ^24^. To further increase biomass we increased input stool weight, optimized volumes of certain key reagents, and processed multiple tubes of stool in parallel for each sample (see methods for details). The final IgA MIG-Seq protocol generated sufficient biomass in the IgA+ microbial fraction to enable deep shotgun metagenomic sequencing (mean extracted DNA of IgA+ fraction = 5.1 ± 0.6 ng) **(Supplemental Table S1)**.

**Figure 1.**
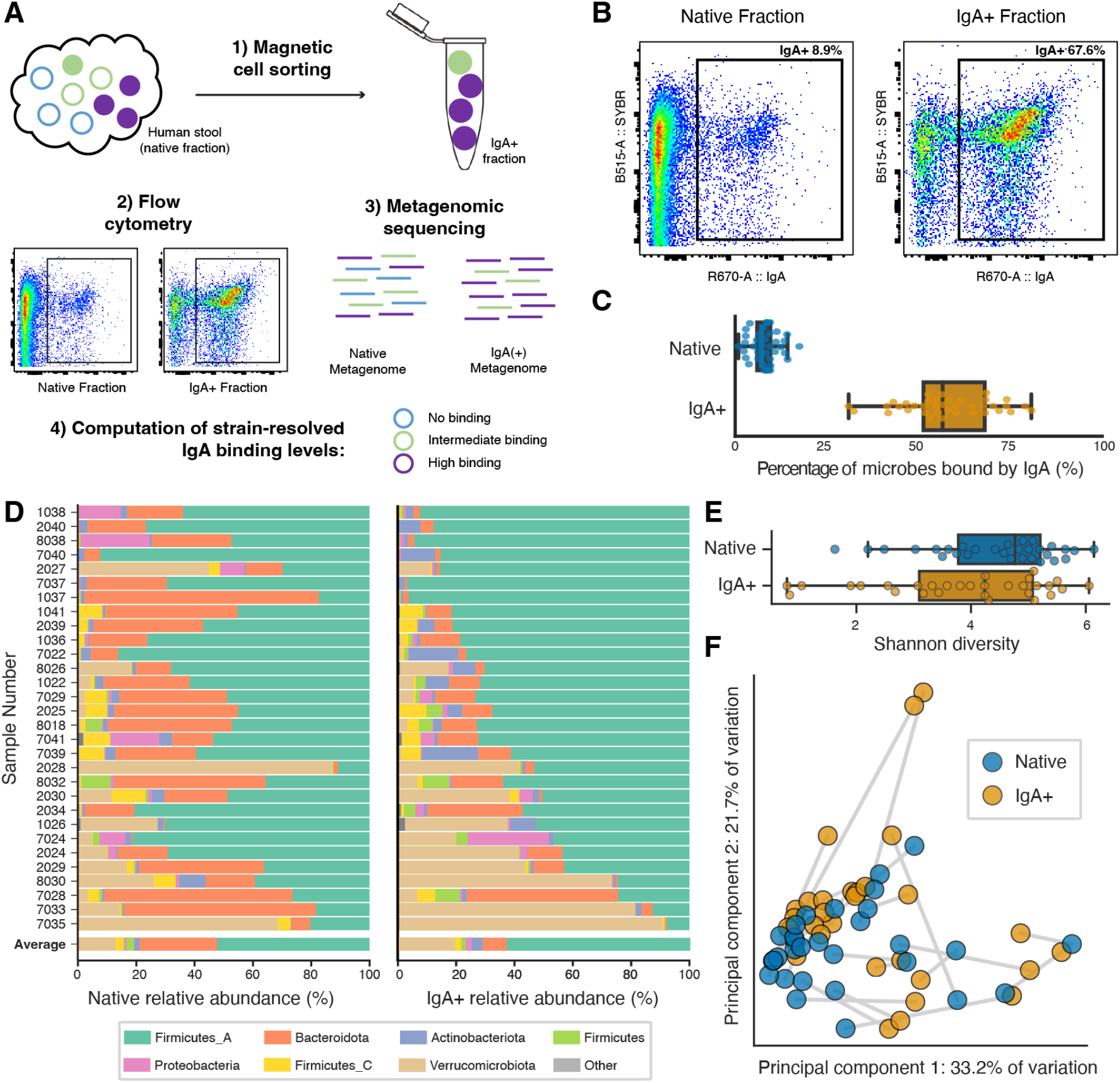
Metagenomic Immunoglobulin Sequencing (MIG-Seq) is a useful and robust technique. **(A)** Schematic overview of the IgA based Metagenomic Immunoglobulin Sequencing workflow. Colors represent different microbial species, and filled circles represent microbes bound by IgA. **(B)** Representative flow cytometry plot of native and IgA+ fractions sorted by magnetic cell separation. **(C)** Percentage of total microbial cells bound by IgA, as measured by flow cytometry, in native samples and MACS sorted IgA+ fractions. **(D)** Phylum-level relative abundance of native metagenomes (left) and IgA+ fraction metagenomes (right) for the 30 samples used in this study. **(E)** Shannon diversity of native (top) and IgA+ (bottom) metagenomes. **(F)** Principal component analysis of weighted UniFrac distance between all metagenomes in this study. Lines connect native and IgA+ metagenomes from the same stool sample.

We next applied IgA MIG-Seq to a set of human adult stool samples that were previously collected during the course of a dietary intervention ^25^. We performed IgA MIG-Seq on 38 total stool samples and thus sequenced 76 metagenomes (one native fraction and one IgA+ fraction for each sample) (mean sequencing depth = 16.2 ± 0.4 giga-base pairs). Five of the 38 stool samples (13%) failed flow cytometry quality control **(Supplemental Table S1)**, and 3 of the remaining 33 stool samples (9%) had either the native or IgA+ fraction below the minimum sequencing depth of 12 Gbp **(Supplemental Table S1)**. We thus conducted our analysis on the 30 fecal samples passing QC and with sufficient sequencing depth. These 30 samples came from 19 different subjects, 11 of which had 2 samples collected (one before and one at the end of the dietary intervention). As outlined in the IgA MIG-Seq protocol, the native metagenomes in this study were collected after resuspending microbial cells in PBS and were processed similarly to the IgA+ samples. This ensured that all differences between the two samples were the result of IgA sorting, and not technical variation to differences in DNA extraction, library preparation, or DNA sequencing protocols **(Supplemental Note S1)**.

The same general IgA binding patterns were observed among the 30 samples analyzed in this study (**Figure 1D; Supplemental Table S2)**. As previously reported ^26–28^, Proteobacteria were significantly more abundant in the IgA+ fraction, and Bacteroidota were significantly depleted in the IgA+ fraction (p = 2.8 x 10^-11^, 6.8 x 10^-40^, n = 102, 492 detected genomes, respectively; Wilcoxon rank-sum test with Benjamini/Hochberg FDR correction). IgA+ fractions had lower Shannon diversity than native samples (**Figure 1E**), as expected given that the IgA+ microbiome is a subset of the native microbiome (*p* = 0.052, n=30, Wilcoxon signed-rank test). While the samples were collected as part of a dietary intervention, we found no significant impact of the dietary interventions on IgA binding levels **(Supplemental Figure S2A)** or microbiome composition in the native or IgA+ fractions (**Figure 1F, Supplemental Figure S2B)**. Finally, IgA+ fractions were found to be significantly more variable on a subject-to-subject level than native samples **(Supplemental Figure S2B**; p=6.7 x 10^-12^). Thus, while there is general conservation between subjects in the taxa that are bound by IgA (**Figure 1D**), IgA targeting is more variable between individuals than microbiome composition overall.

### Microbial IgA targets are generally consistent across healthy humans

We next analyzed IgA coating levels of individual strains using the recently-developed IgA+ probability metric ^24^, defined as the likelihood of IgA binding for a particular taxon. When we created a ranked distribution of all strain-level IgA+ probability values calculated in this study (n = 3,519), three distinct divisions emerged (**Figure 2A**). We classified these groups as “uncoated” (IgA+ probability < 0.005; 21% of species detections), “low binding” (1 < IgA+ probability < 0.005; 61% of species detections), and “high binding” (IgA+ probability > 1; 18% of species detections). Proteobacteria and Firmicutes species were significantly enriched in species with high IgA binding, while Bacteroidota was enriched in uncoated species (**Figure 2B**) (p < 0.01; Fisher’s exact test with Benjamini/Hochberg FDR correction). We identified 20 genera with significantly higher IgA+ probability values than other taxa, many of which belong to *Oscillospiraceae* (a taxonomic family including several species that have been positively associated with human health) (p < 0.01; Mann-Whitney U test with Benjamini/Hochberg FDR correction) **(Supplemental Table S3)**. To explore the hypothesis that IgA may be coating beneficial strains, we examined strains that are considered potentially health-associated. Specifically, we grouped microbes from four genera identified as primary candidates for next generation probiotics (*Bifidobacterium*, *Faecalibacterium*, *Akkermansia*, and *Eubacterium*)^29,30^ as “probionts”. We found these probionts to be more frequently bound by IgA than non-probiont taxa (**Figure 2C**; p = 1.3 x 10^-4^; Wilcoxon rank-sum test). This finding is consistent with previous reports ^18^ and suggests that probionts may occupy a niche in the gut with higher levels of host / microbe immune interactions.

**Figure 2.**
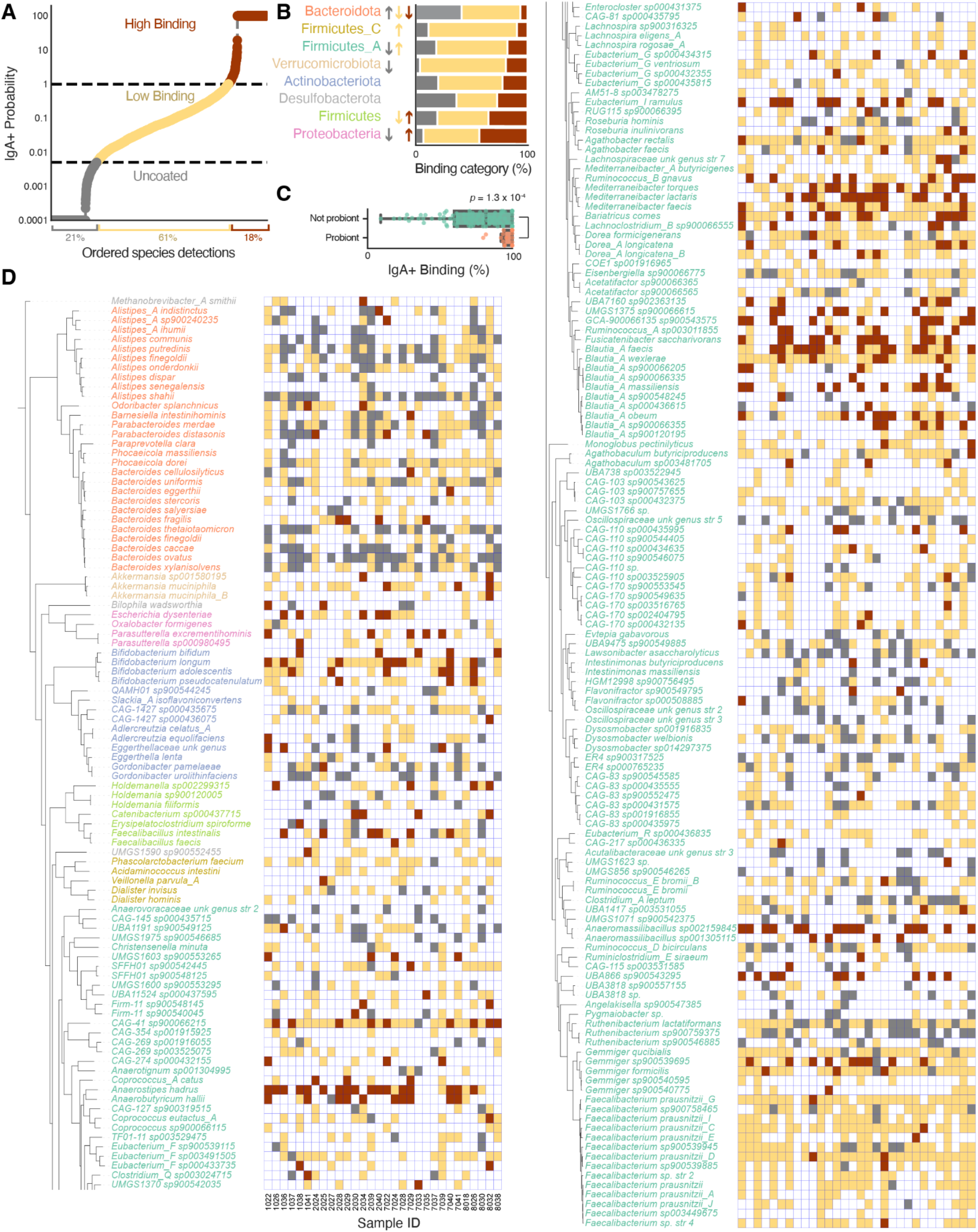
IgA binding levels are consistent across healthy humans. **(A)** Log-scaled IgA+ probability of all species detected in all samples, sorted by rank. Thresholds separating IgA+ probability values classified as “uncoated”, “low”, and “high” are marked with vertical dotted lines at 0.005 and 1. Strains only detected in the IgA+ or native fractions were assigned IgA+ probability values of 100 and 0.0001, respectively. **(B)** The percentage of detections within each phylum displaying the various binding levels. Colors match those in subpanel A. Arrows indicate phyla that are significantly enriched (upward facing arrow) or depleted (downward facing arrow) in a particular binding category (color of arrow) (*p* < 0.05; Fisher’s exact test with Benjamini/Hochberg FDR correction). **(C)** The percentage of detections with high or low IgA binding among probiont and non-probiont genera. P-value from Wilcoxon rank-sum test. **(D)** Concatenated gene phylogenetic tree of all species detected in at least 5 samples with IgA coating across samples. Heatmap colors match those in subpanel A, and unfilled boxes indicated that the species was not detected in that sample.

To examine IgA binding at the species level, we generated a phylogenetic tree of all microbes detected in at least 5 samples and overlaid IgA binding data (**Figure 2D**). Species-level IgA binding patterns were remarkably consistent across the individuals and samples in this study, demonstrating overall conservation of microbial targeting by human IgA (**Figure 2D**). While clear trends are observed in line with the phylogenetic conservation mentioned earlier (e.g., generally uncoated Bacteroidota strains and generally high coating of Proteobacteria strains), we also found that certain strains exhibited unique binding patterns that diverged from their phylogenetic group. For example, *Bacteroides fragilis* exhibited high levels of IgA binding even though it belongs to the phylum with the lowest levels of IgA binding overall (Bacteroidota) (**Figure 2D; Supplemental Figure S3)**. The detection of species with patterns of IgA coating distinct from their phylogeny suggests unique genetic content within strains that drives IgA binding.

### Specific microbial genes are associated with increased IgA binding

We next leveraged the whole-genome sequencing data provided by IgA MIG-Seq to examine connections between microbial genes and IgA binding. A random forest machine learning classifier trained to predict microbial IgA binding levels based on microbial gene content exhibited 75% accuracy overall (**Figure 3A**). The model was more accurate in classifying “uncoated” and “high binding” than “low binding” species (0.81, 0.86, and 0.68 area under curve (a.u.c), respectively), as “high binding” and “uncoated” microbes were more often mis-classified as “low binding” than as each other (**Figure 3B**). These data highlight both the robust association between microbial gene content and IgA binding levels and the hierarchical nature of binding levels (e.g. highly-bound microbes have more similar gene content to lowly-bound microbes than uncoated microbes).

**Figure 3.**
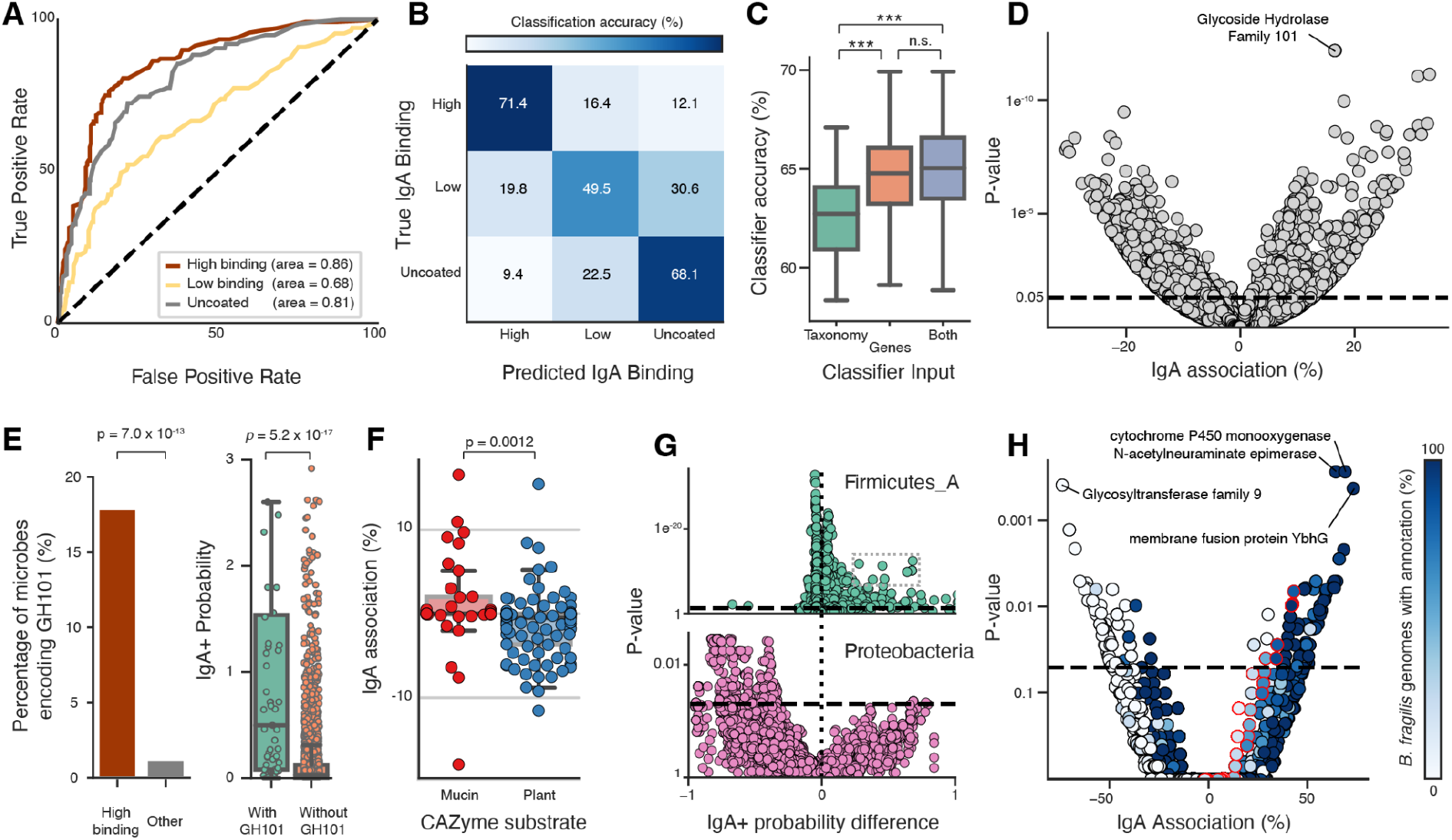
Taxon-specific microbial genes are associated with high IgA targeting. **(A)** Receiver Operating Characteristic (ROC) Curve displaying the performance of the random forest classifier in classifying a microbe’s IgA binding class based on gene content. The area under the curve (AUC) quantifies the overall ability of the classifier to distinguish between each class versus the other classes. **(B)** Confusion Matrix depicting the true (y-axis) versus predicted (x-axis) classifications of microbial samples’ IgA binding classes. **(C)** One hundred random forest classifiers each were trained on microbial taxonomy, microbial gene content, or both. Accuracy values were compared using the Wilcoxon rank-sum test. *** = *p* < 1E-5; n.s. = not significant. **(D)** For each microbial function (dot), the association of that function with IgA binding (x-axis; percentage of microbes with high IgA coating that encode this function - percentage of microbes with high IgA coating that do not encode this function) versus the p-value of the association of the function with high IgA coating (Fisher’s exact test with FDR correction). Horizontal dotted line drawn at p=0.05. **(E)** Barplot comparing the total percentage of microbes with high IgA coating that encode the GH101 versus that of lowly coated or uncoated microbes. P-value from Fisher’s exact test (left). Boxplot comparing the IgA+ probability of microbes that encode GH101 versus the IgA+ probability of lowly coated or uncoated microbes. P-value from Wilcoxon rank-sum test (right). **(F)** For each CAzyme (dot), the association of that gene with IgA binding (see subpanel D x-axis for description) with predicted CAzyme substrate. P-values calculated using Wilcoxon rank-sum test. **(G)** For each microbial function (dot), the association of that function with IgA+ probability (x-axis; median IgA+ probability of microbes encoding function - median IgA+ probability of microbes not encoding function) versus the p-value of the association of the function with IgA+ probability (Wilcoxon rank-sum test with FDR correction). Horizontal dotted line at p=0.05. **(H)** Same as (D) but only considering species of *Bacteroides*. Color depicts the fraction of *Bacteroides fragilis* genomes that encode each annotation. Dots with red outline are exclusively detected in *B. fragilis*.

To determine the relative significance of microbial gene content versus taxonomy in relation to IgA binding, we trained additional machine learning classifiers based on i) taxonomy, ii) gene content, or iii) both taxonomy and gene content. Classifiers based on gene content were more accurate than those based on taxonomy (*p* = 6.2E-9; Wilcoxon rank-sum test), but classifiers with access to both taxonomy and gene content performed no better than those with gene content alone (**Figure 3C**). Two hypotheses emerge from these findings. First, given that microbial gene content is more predictive of IgA binding than taxonomy, we hypothesize that certain genes regulate a microbe’s IgA binding independent of taxonomy. Second, since taxonomy can moderately predict IgA binding on its own, but does not add additional accuracy on top of gene content, we hypothesize that some genes associated with IgA binding display strong taxonomic linkage. Evidence supporting both of these hypotheses is presented below.

When considering all microbes detected across all samples in this study, glycoside hydrolase family 101 (GH101) emerged as the annotation most significantly associated with high IgA binding (**Figure 3D, E)**. GH101 family members are endo-glycoside hydrolases involved in mucus degradation via the liberation of glycans from host mucin proteins ^31^. We identified GH101 in 10 genomes distributed across 3 phyla and 8 genera, including the species *Bifidobacterium longum*, *Mediterraneibacter torques*, and *Odoribacter sp900544025*. This gene’s wide taxonomic distribution, strong association with IgA binding, and known function associated with host mucus degradation, confirms the existence of microbial functions that increase IgA binding regardless of taxonomy. Additionally, our results show that CAZymes involved in mucus degradation are generally more associated with high IgA binding compared to CAZymes predicted to act on other substrates (**Figure 3F**).

A set of 10 annotations with a strong and significant IgA association were also identified among Firmicutes_A (**Figure 3G**). Similar to GH101, their functions primarily involve enzymes and domains associated with the breakdown or interaction with mucin and mucus components **(Supplemental Table S4)**. Interestingly, examination of other phyla for the relationship between IgA coating and enriched functions presented different plot structures, possibly indicating distinct phylum-level IgA associations **(Supplemental Figure S4)**. Proteobacteria, for instance, displayed far more annotations that were anti-associated (n=350) than positively associated (n=7) with IgA binding (**Figure 3G**), potentially suggesting that IgA binding is the “norm” in this phylum and that certain genes and/or taxa have adapted to escape binding. Among all genera, *Bacteroides* had the most annotations significantly associated with high IgA binding (n=150; **Figure 3H; Supplemental Table S4**). As *Bacteroides fragilis* exhibited substantially higher IgA binding than other *Bacteroides* (**Figure 2D**), we next investigated whether this species, in particular, was driving the high-IgA annotations detected. A clear link was observed between 1) annotations associated with IgA binding among all *Bacteroides*, and 2) annotations associated with the species *Bacteroides fragilis* in particular (**Figure 3H**). This finding demonstrates the existence of genes strongly linked to both IgA binding and taxonomy.

### IgA binding reduces *in situ* microbial replication rates

We next used the shotgun metagenomic sequencing data generated by IgA MIG-Seq to test the “enchained growth” model, which proposes that polyreactive IgA might preferentially bind bacteria with high replication rates ^19^. We calculated the *in situ* microbial replication rates, measured as iRep values ^23^, of all microbes detected in this study. We controlled for strain-specific properties that can impact iRep ^23^ by only comparing values from the same strain detected in the IgA+ versus native fractions of the same stool sample. Among species with low or high IgA coating, replication rates were significantly higher in the native compared to the IgA+ fractions (**Figure 4A; Supplemental Table S5**). Significant variation was also observed across microbial phyla in the magnitude of replication rate reduction **(Supplemental Figure S5)** (p = 0.01; Kruskal-Wallis test), with Bacteroidota exhibiting a greater decrease in iRep in native versus IgA+ samples than Firmicutes and Firmicutes_A. These data support the ability of IgA to influence bacterial replication rates, provide empirical evidence supporting the “enchained growth” model, and highlight the varying impacts of IgA binding across taxonomic groups.

**Figure 4.**
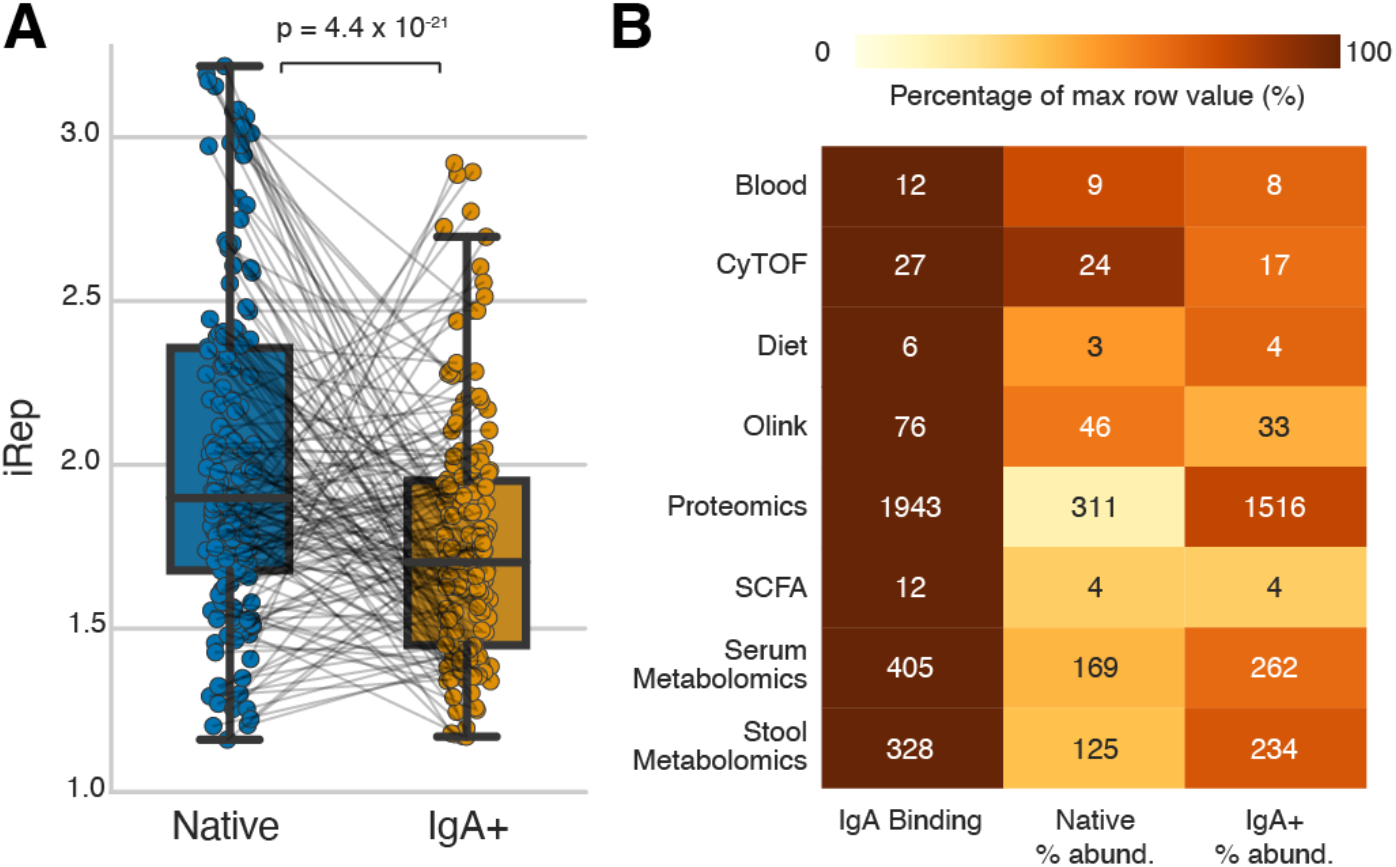
IgA binding impacts *in situ* microbial replication rates and is integrated with other components of the immune system. **(A)** Paired comparison of the replication rates (iRep) of species with low or high IgA binding in native (left) and IgA+ (right) fraction of the same fecal sample (paired values connected with lines). P-value from Wilcoxon signed-rank test. **(B)** A phylum-level correlation analysis between 3 distinct microbial measurements (x-axis) and 9 measurements of human health and microbiome function (y-axis). Numbers in cells indicate the number of significant (p < 0.05) correlations identified after FDR correction. Colors represent the percentage of significant associations between each health metric and microbial metric, normalized to the maximum value in each row (highlighting the microbiome metric with the most significant associations with each immune metric).

### IgA coating is tightly associated with other components of the immune system

Comprehensive high-resolution, multi-omic immune and health data were previously generated from the same subjects and stool samples analyzed in this manuscript ^25^, allowing us to investigate the relationship between IgA binding and human health status. Available data includes blood markers like triglyceride and blood glucose levels (n=6), immune cell frequencies as quantified by CyTOF (n=17), detailed dietary information (n=18), an Olink panel of inflammation-linked cytokines (n=67), stool short chain fatty acid quantification (n=9), untargeted stool proteomics (n=5,682), untargeted stool metabolomics (n=479), and untargeted serum metabolomics (n=321). We observe many correlations between the immune and health data with both the native and IgA+ relative abundance values (**Figure 4B, right two columns; Supplemental Table S6)**. However, the ratio of these two measures, IgA+ relative abundance to native relative abundance, demonstrated much stronger correlations with the immune and health data (**Figure 4B, left column; Supplemental Table S6).** The fact that the combined assessment of these two measurements holds greater informational value in relation to diverse other measures of the host system than considering them independently is particularly striking given the prevalent usage of native relative abundance as a key microbiome metric. Together these data reinforce IgA binding as a highly relevant measurement in the context of microbiome-immune system interactions.

A number of compelling associations emerged from the analysis with host-derived data (**Table 1) (Supplemental Table S6).** Triglyceride levels were significantly associated with IgA+ binding of the Firmicutes_A, Actinobacteriota, and Bacteroidota phyla, potentially reflecting the impact of systemic inflammatory responses or metabolic homeostasis on IgA targeting. Firmicutes_A IgA binding was highly correlated with Central Memory CD8 T Cells and anti-correlated with fiber intake, supporting the role of diet in shaping the IgA response against specific microbes. The association between Firmicutes IgA binding and valeric acid levels, a key SCFA known for its anti-inflammatory properties, highlights the crosstalk between the host’s immune response, the microbiota, and the metabolic landscape of the gut. Collectively, these results illustrate how IgA targeting is integrated with overall physiology and health status in humans and that this novel assay, MIG-Seq, is a powerful new tool to reveal mechanisms of this integrated system.

**Table 1.** Most significant correlations between phyla IgA binding and immune metrics. Variables negatively associated with IgA binding are in bold.

| Measurement | Phylum | Variable | p-value | Rho |
| --- | --- | --- | --- | --- |
| Blood | Firmicutes_A | Triglycerides | 1.4E-06 | 0.14 |
|  | Actinobacteriota | Triglycerides | 1.9E-04 | 0.37 |
|  | Bacteroidota | Triglycerides | 9.7E-04 | 0.23 |
|  | Proteobacteria | LDL Cholesterol | 1.5E-03 | 0.46 |
|  | Verrucomicrobiota | Insulin | 3.1E-02 | 0.61 |
| CyTOF | Firmicutes_A | CM CD8 T Cells | 6.2E-14 | 0.22 |
|  | Actinobacteriota | CD8 T Cells | 4.1E-04 | 0.40 |
|  | Bacteroidota | pDCs | 4.1E-04 | 0.26 |
|  | Firmicutes | CM CD8 T Cells | 5.1E-04 | 0.44 |
|  | Proteobacteria | mDCs | 8.8E-03 | 0.42 |
| Diet | **Firmicutes_A** | **Fiber (Other)** | **6.0E-05** | **-0.13** |
|  | **Bacteroidota** | **Total Fiber** | **5.1E-04** | **-0.20** |
|  | **Proteobacteria** | **Yogurt** | **4.5E-02** | **-0.38** |
| Olink | Firmicutes_A | MMP-10 | 1.8E-13 | 0.21 |
|  | Actinobacteriota | CCL4 | 8.3E-08 | 0.50 |
|  | **Firmicutes** | **CX3CL1** | **2.9E-03** | **-0.40** |
|  | Proteobacteria | LAP TGF-beta-1 | 3.6E-03 | 0.47 |
|  | Bacteroidota | CST5 | 8.2E-03 | 0.21 |
| Proteomics | **Firmicutes_A** | **CUU 3304** | **4.7E-24** | **-0.29** |
|  | **Actinobacteriota** | **BACDOR 00299** | **8.7E-10** | **-0.56** |
|  | **Bacteroidota** | **RO1 21290** | **2.2E-06** | **-0.32** |
|  | **Firmicutes** | **P61626** | **1.2E-03** | **-0.52** |
|  | **Verrucomicrobiota** | **HMPREF0322 02320** | **1.0E-02** | **-0.84** |
| SCFA | Firmicutes_A | Isobutyric Acid | 2.2E-04 | 0.11 |
|  | Actinobacteriota | Lactic Acid | 6.9E-03 | 0.28 |
|  | Firmicutes | Valeric Acid | 2.0E-02 | 0.31 |
|  | Bacteroidota | Isobutyric Acid | 2.0E-02 | 0.16 |
| Serum Metabolomics | Firmicutes_A | Asparagine | 4.3E-14 | 0.24 |
|  | Actinobacteriota | Propionylcarnitine | 1.5E-08 | 0.56 |
|  | **Bacteroidota** | **1,5,6,7-Tetrahydro-4H-Indol-4-One** | **6.4E-06** | **-0.35** |
|  | **Firmicutes** | **Omeprazole** | **1.1E-05** | **-0.64** |
|  | Proteobacteria | Deoxycarnitine | 1.1E-02 | 0.49 |
| Stool Metabolomics | **Firmicutes_A** | **2'-Deoxycytidine 5'-Monophosphate** | **8.5E-18** | **-0.25** |
|  | **Actinobacteriota** | **Pipecolic Acid** | **1.2E-06** | **-0.48** |
|  | **Bacteroidota** | **Kynurenine** | **2.1E-05** | **-0.31** |
|  | **Firmicutes** | **Methylsuccinic Acid** | **1.1E-03** | **-0.52** |
|  | Proteobacteria | Heptanoic Acid | 1.4E-02 | 0.53 |

## Discussion

Here, we present a technique to measure IgA coating of the human intestinal microbiota with strain-level resolution. By applying this method to a set of fecal samples from healthy adults, we demonstrate its clear utility over previous IgA measurement techniques, and well as its complementarity to commonly-used microbial relative abundance measures. Specifically, the strain-level resolution allowed us to analyze specific IgA targeting patterns among bacteria and archaea (**Figure 2**), identify microbial genes associated with IgA targeting (**Figure 3**), and measure the *in situ* growth rates of bacteria coated with IgA (**Figure 4**). None of these are possible with IgA-SEQ^TM^ due to its reliance on 16S sequencing. Furthermore, finding that strain-level IgA binding values are significantly more associated with human immune and dietary status than microbial relative abundance (**Figure 5**) demonstrates the exciting potential of IgA MIG-Seq to identify new human host-microbiome interactions that would be missed by today’s methods. The innovation of IgA MIG-Seq lies in both the protocol to generate sufficient biomass for modern deep shotgun sequencing, and the development and utilization of bioinformatics algorithms to analyze the data in a robust manner (see **Supplemental Note S2** for discussion of MIG-Seq compared to previously-developed IgA sequencing techniques).

We identified many of the same taxonomic IgA patterns that have been previously reported ^26–28^ (**Figure 2**), and we provide new compelling evidence that gene content is undergirding these taxonomic associations (**Figure 3**). We thus argue that focusing on microbial genes, rather than on taxonomy, will lead to more insight into IgA dynamics. The microbial gene most associated with IgA binding in our study is GH101, a mucus degradation gene found across many taxonomic groups (**Figure 3**). We do not suggest that the physical GH101 proteins are targeted by IgA (as IgA tends to bind cell surface antigens ^11^); rather, we hypothesize that mucus utilizing bacteria are targeted by IgA due to their proximity to the host and increased interaction with the immune system. This niche is not typically associated with enteric pathogens ^32^, but rather is associated with probionts like *Bacteroides* and *Akkermansia muciniphila.* This synergizes with our finding of increased IgA coating of probiont species (**Figure 2C**).

As posited by the adhesion-based IgA model ^18^, probionts may be actively encouraged to colonize the mucus via host IgA targeting. *Bacteroides fragilis* is the canonical species that has been experimentally shown to benefit from IgA coating via a mucus-anchoring mechanism ^17^, and in this study we report uncharacteristically high IgA binding levels of *B. fragilis* (**Figure 2D**). It is likely that microbes colonizing the mucus layer have evolved strategies to mitigate and tolerate host immune recognition, and the additional genetic content associated with IgA coating may represent genetic content gained to directly promote immune tolerance and a peaceful co-existence with the host while occupying this host-associated niche. A key example of this is a lipid produced by *Akkermansia muciniphila* that promotes homeostatic immunity and may have therapeutic potential as an anti-inflammatory agent ^33^. Thus, IgA coating may prove to be a useful metric by which to identify promising candidates for next-generation probiotics that modulate the immune system, and MIG-Seq can be used to identify specific genetic content or subsets of strains within species that could be leveraged for therapeutic potential.

The relationship between humans and our microbiota, which has critical roles in human organismal homeostasis, has co-evolved so that our immune system is endowed with the ability to shape the microbiome, and the microbiome is conversely endowed with potent immune-modulating properties. Mucosal IgA appears to be a key effector module of this interaction due to its remarkably high production levels, potential for precise specificity, and the ability to cause both beneficial and deleterious fitness impacts. It follows that the strains targeted by IgA are actively recognized by the immune system, as indicated by our findings, and thus worthy of special consideration. The importance of IgA is in stark contrast to our limited understanding of its mechanistic interactions with our microbiota. This knowledge gap can likely be attributed to the only recent development of high-throughput techniques to holistically profile IgA binding, coupled with the significant limitations associated with 16S-based methods. The metagenomic IgA sequencing protocol presented here will enable future studies to identify microbial IgA coating levels with unprecedented levels of resolution, and catalyze a shift in our understanding of host-microbiome interactions.

## Acknowledgements

We would like to thank Drs. Lilian H. Lam, Holden Maeker, Jessica Fessler, Tadashi Takeuchi, Matthew Carter, and KC Huang for helpful discussions during the preparation of this manuscript.

Support for the project was provided from National Institutes of Health grant F32DK128865 (M.R.O.), National Institutes of Health training grant T32 AI007328-30 (M.R.O.), the Colleen and Robert D. Hass fund (S.P.S.), and National Institutes of Health grants T32DK007056, K08DK134856 (S.P.S.), R01DK085025 (J.L.S.), and DP1AT009892 (J.L.S.). The content is solely the responsibility of the authors and does not necessarily represent the official views of the National Institutes of Health. J.L.S. is a Chan Zuckerberg Biohub Investigator.

This publication includes data generated at the UC San Diego IGM Genomics Center utilizing an Illumina NovaSeq 6000 that was purchased with funding from a National Institutes of Health SIG grant (#S10 OD026929). This publication includes data generated at the Stanford Shared FACS Facility (NIH S10 Shared Instrument Grant 1S10OD026831-01).

## Data availability

The data supporting the findings of this study are available within the paper and its supplementary information files. Metagenomic reads are in the process of being submitted to the SRA and will be made freely available.

## Contributions

M.R.O, S.P.S, and J.L.S. designed the study. S.P.S. led development of MIG-Seq protocol. M.R.O. performed bioinformatic analysis. M.R.O, S.P.S, and E.L. performed wet lab experiments. M.R.O. wrote the manuscript and all authors contributed to manuscript revisions.

## Supplemental Tables

**Supplemental Table S1**. Information on the DNA extraction concentration, sequencing depth, IgA sorting information, and metadata for all 38 samples.

**Supplemental Table S2**. Microbial relative abundance and IgA+ probabilities for all samples.

**Supplemental Table S3**. Statistics for IgA binding enrichment among phyla and genus level taxonomic groups.

**Supplemental Table S4.** Statistics for association of microbial annotations with IgA binding.

**Supplemental Table S5.** Paired iRep values for all species detected in both IgA+ and native samples.

**Supplemental Table S6.** Statistical associations between each phylum and each immune metric.

## Supplemental Figures

**Supplemental Figure S1.**
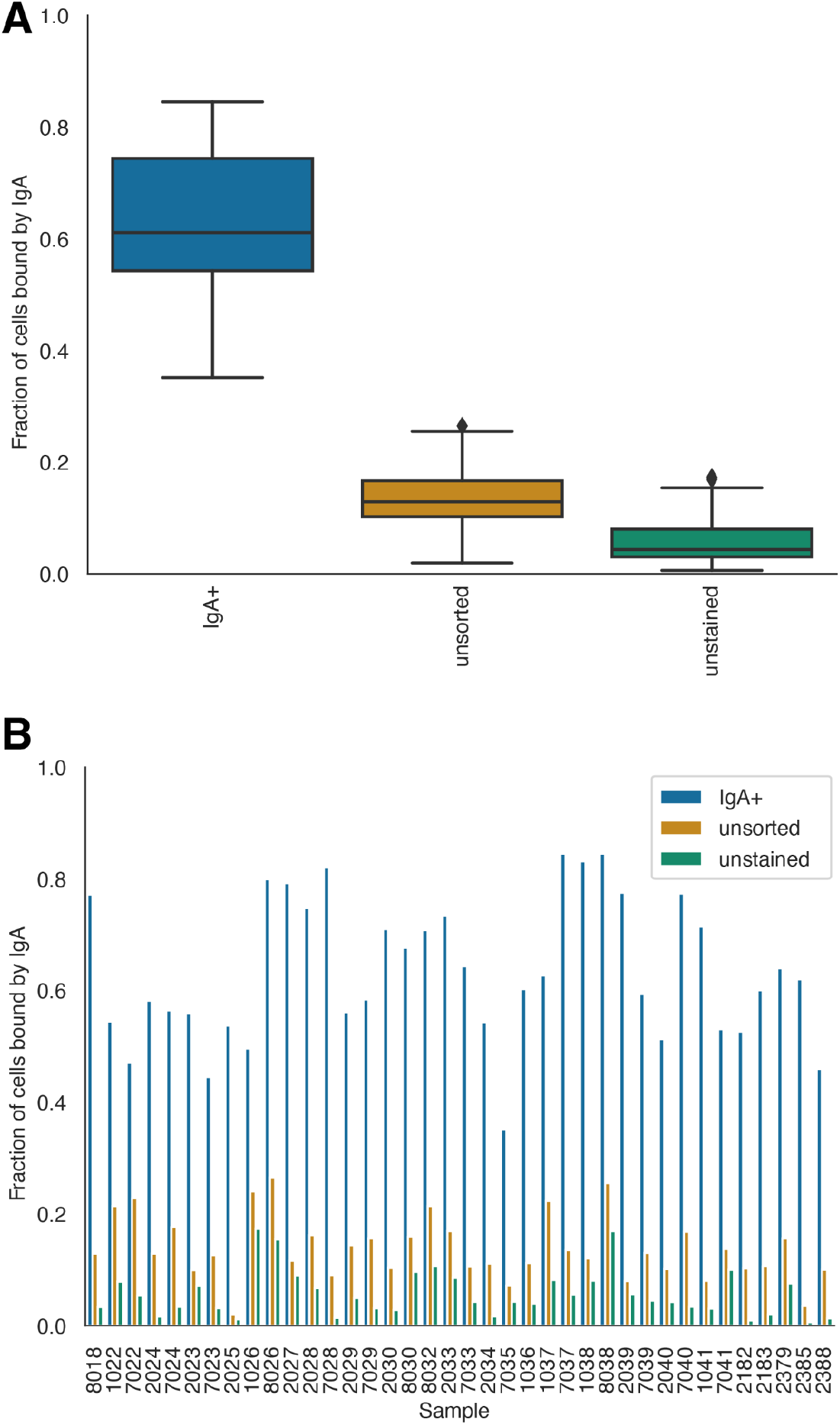
Effectiveness of magnetic cell sorting for enriching IgA+ cells. Aggregated **(A)** and sample-level **(B)** fraction of bacterial cells bound by IgA as assessed by bacterial flow cytometry. IgA+ (blue) is the result of magnetic cell sorting, unsorted (orange) is the native unsorted sample, and unstained (green) is the unsorted fraction without SYBR dye added.

**Supplemental Figure S2.**
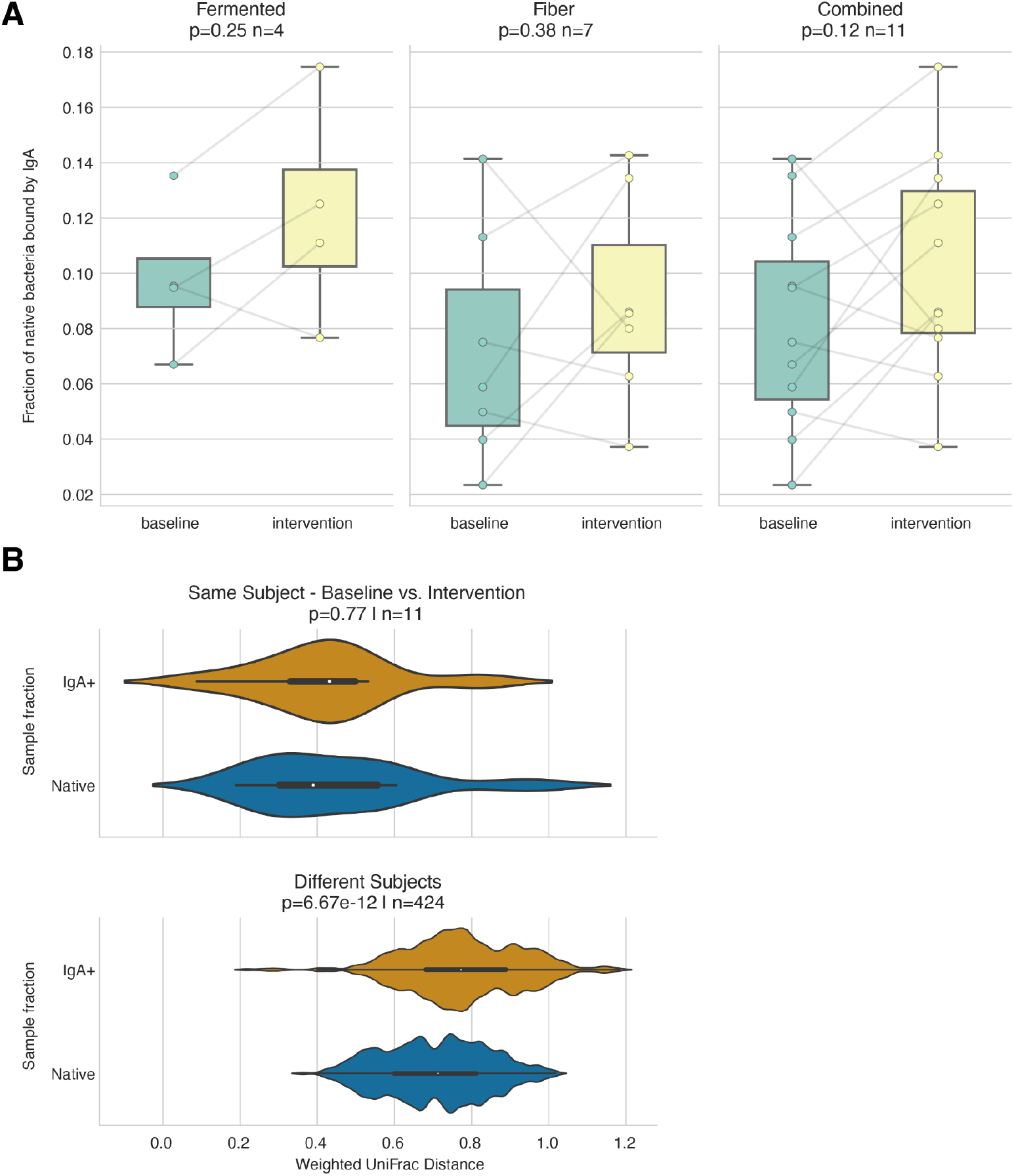
**A)** Boxplots comparing the overall fraction of bacteria bound by IgA in native samples (as assessed by bacterial flow cytometry) in baseline samples versus after dietary intervention of increased consumption of fermented foods (left) or fiber (middle). The impact of dietary interventions was also assessed in aggregate (right; a combination of the other 2 plots). Lines connect samples from the same subjects over time. P-value from Wilcoxon signed-rank test. **B)** Boxplot of weighted UniFrac distance between the same subjects in baseline vs. intervention samples (top) and weighted UniFrac distance between different subjects (bottom). P-values from comparing the distribution of IgA+ vs native weighted UniFrac distances using Wilcoxon rank-sum test.

**Supplemental Figure S3.**
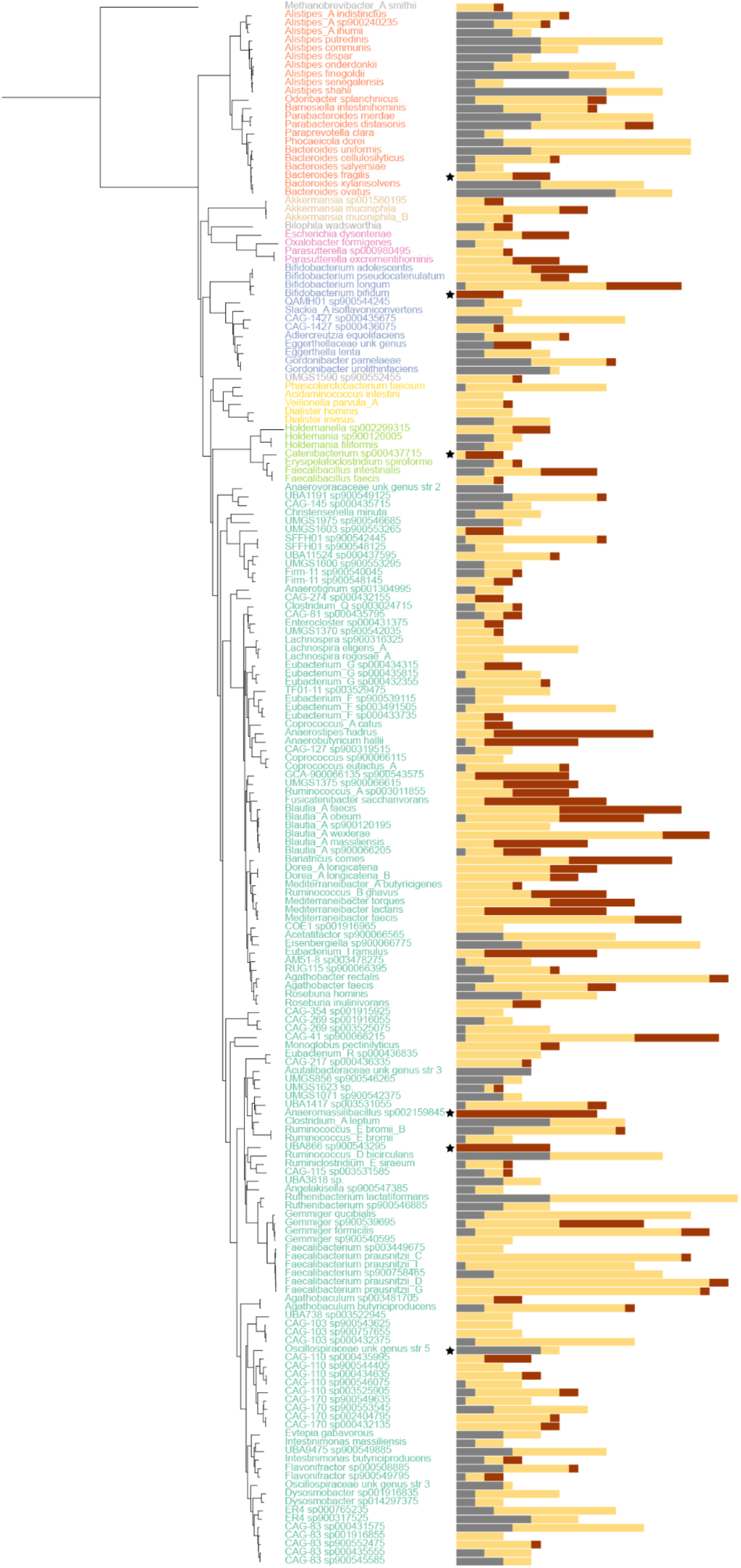
A concatenated gene phylogenetic tree of all microbial strains detected in at least 5 samples. Stacked barplots on the right display the relative abundance of IgA binding categories for each detection. The total bar length represents the total number of detections. Black stars mark taxa that appear to have substantially different IgA coating than their phylogenetic neighbors based on manual inspection.

**Supplemental Figure S4.**
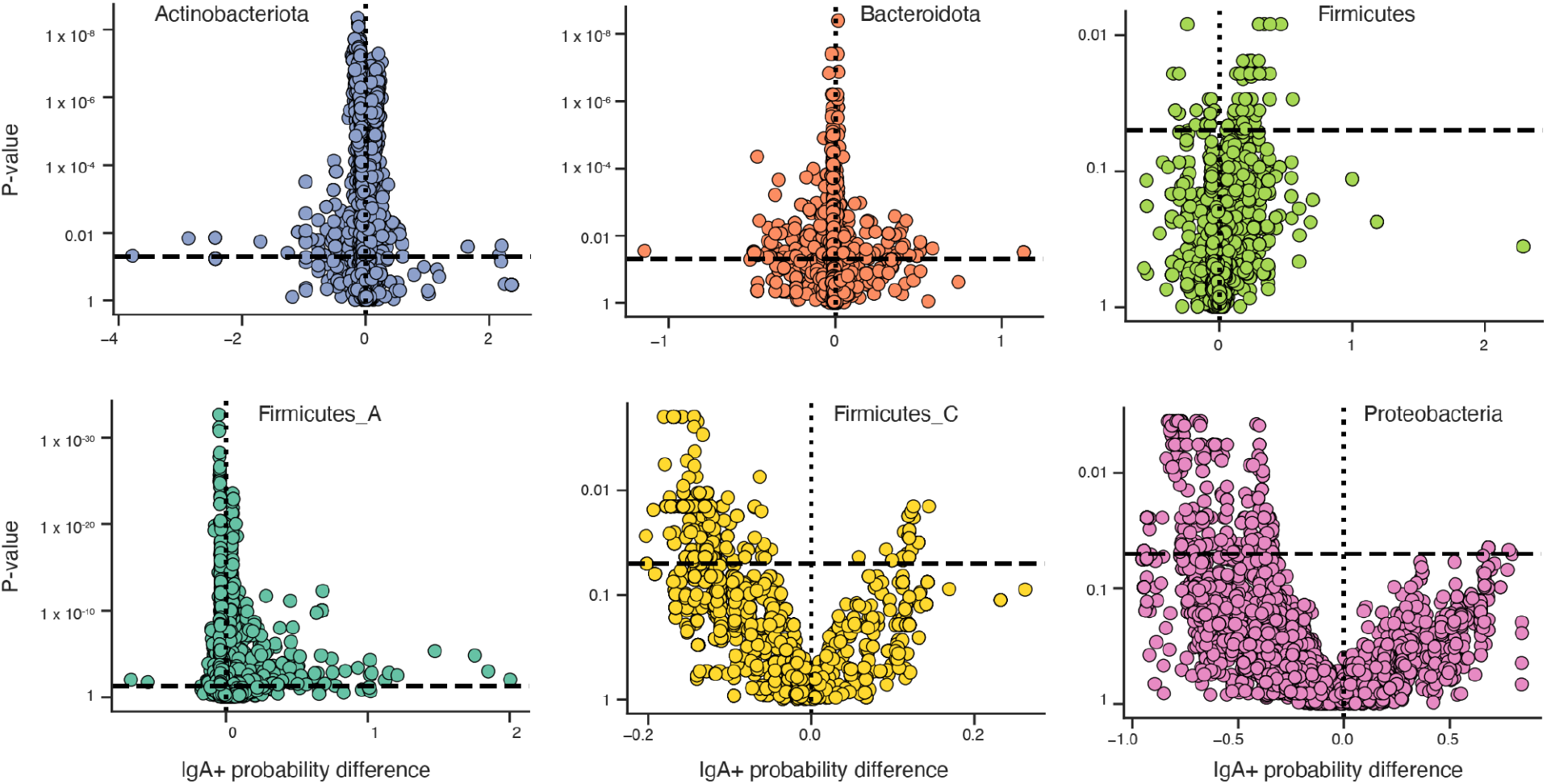
For each microbial function (dot), the association of that function with IgA+ probability (x-axis; median IgA+ probability of microbes encoding function - median IgA+ probability of microbes not encoding function) versus the p-value of the association of the function with IgA+ probability (Wilcoxon rank-sum test with FDR correction). Horizontal dotted line at p = 0.05. Each phylum is treated independently.

**Supplemental Figure S5.**
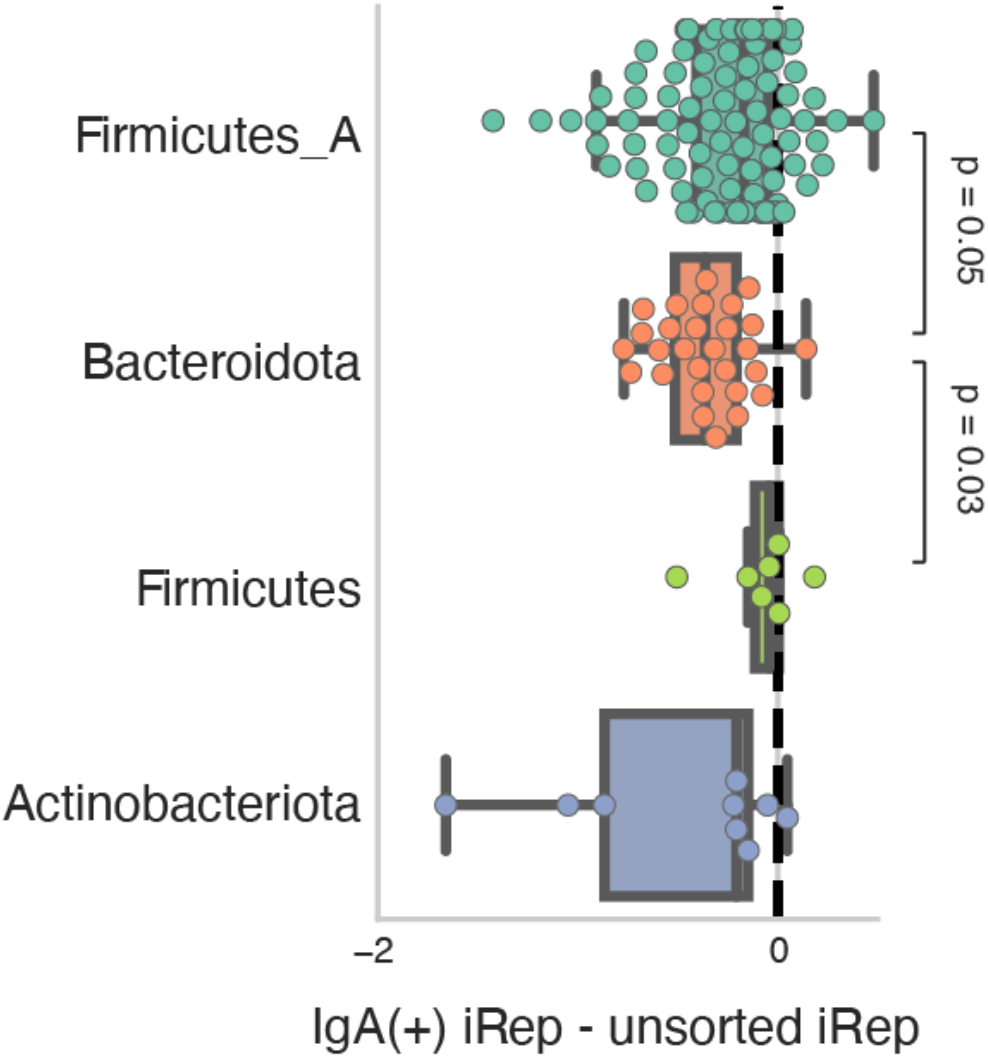
For each phylum, the difference in iRep in IgA+ vs. native fractions of all species detected within that phylum. Negative values indicate IgA is associated with reduced replication rates. P-values from post hoc pairwise test for multiple comparisons of mean rank sums (Dunn’s test).

## Methods

### IgA Metagenomic Immunoglobulin Sequencing (MIG-Seq)

#### Sample preparation and antibody staining

Frozen human stool samples were thawed on ice; 300 mg of each sample was allotted into a 2 mL tube. 1.25 mL of cold PBS was added and incubated for 5 minutes on ice, followed by vigorous vortexing and pipetting to ensure homogenization of the sample. The sample was then centrifuged at 500 g for 15 min. 1 mL of supernatant was removed and passed through a 70 µm filter. At this point 100 µL (1/10th of sample) was placed on ice and saved as the “native” fraction for subsequent metagenomic sequencing. 450 µL each was aliquoted into two separate wells of a deep-well 96 well plate and treated as individual samples throughout the remainder of the protocol; the two samples are combined at the end of the protocol to increase sample biomass. The plate was centrifuged at 5,000 g for 5 minutes to pellet the sample and supernatant was discarded. Pellet was resuspended with 100 µL of antibody master mix containing (1:30 dilution of Anti-Human IgA, APC [Miltenyi, Cat#130-113-472], 1:30 dilution mouse serum [Jackson labs, Cat #15000120], in a staining buffer [PBS, 3% Fetal Bovine Serum, 0.5% Sodium Azide, 1 mM EDTA]) and incubated for 30 minutes on ice. Samples were washed 1x with cold staining buffer (see above) and centrifuged at 5,000 g for 5 minutes. Supernatant was decanted, and the sample was resuspended in 200 µL of staining buffer. 20 µL of the 200 µL (10%) was removed for a stained, but unsorted control for Flow Cytometry analysis to establish background to calculate “Native IgA binding percentage”.

#### Magnetic cell separation

The remaining 180 µL was stained with 20 µL of anti-APC beads (Stemcell technologies, EasySep APC Positive Selection Kit II, Cat #17681) for 15 minutes at room temperature. 22 µL of magnetic beads were next added (Stemcell technologies, EasySep APC Positive Selection Kit II, Cat #17681) and incubated for 10 minutes at room temperature. Samples were then placed on a 96- well plate magnet for approximately 5 minutes to allow for magnetic separation. The unbound supernatant was carefully discarded. The samples were removed from the magnet, washed with 200 µL of staining buffer, and again magnetically enriched for 5 minutes. This process was repeated for a total of 3, 5-minute magnetic enrichments and washes. After the final enrichment, the duplicate wells (established above) were combined to establish the “IgA positive” fraction.

#### Flow cytometry analysis of samples

Samples were stained with 1x SYBR (Invitrogen, Cat#G9348) in 200 µL of staining buffer. Cells were analyzed on a flow cytometer (BD FACSymphony™) at the Stanford Shared FACS Facility. Fractions analyzed were: a) Native, unstained b) Native, stained with anti-IgA c) IgA+ Fraction after sorting, stained with anti-IgA. For each sample, the native “IgA fraction” was calculated as the number of IgA+ cells in the native sample (b) minus the number of IgA+ cells in the unstained control (a). Samples in which the unstained control (a) had a higher number of IgA+ cells than the native stained sample (b) (4 of 38 samples) or the native stained sample (b) had a higher number of IgA+ cells than the IgA+ sample (c) (1 of 38 samples) were considered to have “failed QC” and were removed from subsequent analysis. Flow cytometry results were visualized using FlowJo™ (BD Life Sciences).

#### Metagenomic Sequencing

DNA extraction, library preparation, and metagenomic sequencing was performed at the UC San Diego IGM Genomics Center. DNA extraction was performed using the Thermo MagMAX Microbiome Ultra kit. DNA quantification was performed using picogreen dye. Roche KAPA HyperPlus and 96-UDI plates were used for library preparation. In this study, 1.75 ng of input DNA and 14 PCR cycles were used during library preparation. A shallow MiSeq Nano sequencing run was used to normalize read sequencing depth. 2 x 150bp reads were then generated on the Illumina NovaSeq 6000 at a target depth of 15 Gbp per sample. See **Supplemental Table S1** for exact sequencing depths per-sample. The metagenomic sequencing reads were trimmed and deduplicated using fastp ^34^. Human reads were removed by mapping reads to the hg17 human genome (GCF_000001405.11) with Bowtie2 ^35^.

### Construction of microbial genome database

To create a study-specific genome database, we used the full set of metagenomes previously generated and published from these stool samples (mean sequencing depth = 7.54 ± 0.38 gigabase pairs) ^25^. Metagenomes were assembled individually, as well as co-assembled with all metagenomes generated from each individual (metaSPAdes; v3.13) ^36^ using unmerged forward/reverse and merged reads (-k 21,33,55,77) with error-correction enabled. Assembly size and contig metrics were evaluated (QUAST73 v5.0) ^37^ and filtered to contigs >=1500 bp for all subsequent analyses. For each assembly, reads from all samples from that individual were mapped (Bowtie2; –very-sensitive -X 1000) and genome bins generated using the contig depth for all samples (MetaBAT278; v2.15, default settings) ^38^. Genome bin quality was assessed using CheckM v1.1.275 ^39^.

Species-level representative genomes were chosen using a series of dRep (v3.0.0) ^40^ commands. First, all genomes binned in this study were de-replicated using the command “dRep dereplicate - sa 0.95 --multiround_primary_clustering --S_algorithm fastANI -comp 50 -con 15”. Next, all representative genomes from this de-replication were compared with all representative genomes in UHGG v1 ^41^ using the command “dRep compare -sa 0.95 --multiround_primary_clustering -- S_algorithm fastANI”. Representative genomes were manually chosen in Python using the default dRep scoring algorithm, with an extra 20 points given to genomes assembled in this study instead of UHGG. Finally, all representative genomes were compared one last time using the command “dRep compare -sa 0.95 --multiround_primary_clustering --S_algorithm fastANI” and representative genomes were again manually chosen as described above. The final genome database included 4659 representative genomes, 901 of which were assembled in this study and 158 of which represent species not found in UHGG. Taxonomic identity of all representative genomes was determined using GTDB-tk (r202) ^42^.

### Microbial relative abundance calculation and sequencing depth normalization

To determine the relative abundance of each microbial species in each sample, each sample’s metagenomic reads were mapped to the microbial genome database described above using Bowtie2 with default settings. The resulting .bam files were analyzed using inStrain profile under default settings ^43^. All samples with less than 12 giga-base pairs of sequencing depth were excluded from analysis. To normalize sequencing depth, a minimum relative abundance threshold was established based on detection of a 4 mega-base pair genome in a sample with exactly 12 giga- base pairs of sequencing depth (relative abundance threshold = 4,000,000 / 12,000,000,000 = 0.00033). In order for a genome to be considered present in a sample, it has to have 1) a genome breadth > 0.5, and 2) a relative abundance above the 0.00033 threshold in either the native or IgA+ metagenome.

### Microbial composition analysis

Stacked bar charts were created in python using matplotlib ^44^. Only genomes considered “present” using the criteria described above were used in stacked barcharts, and relative abundance values were normalized to sum to 100%. Weighted UniFrac distance was calculated between all samples using skbio.diversity (http://scikit-bio.org). In this calculation, microbial relative abundance values were multiplied by 1.25e9 to estimate the number of reads. The tree used in the UniFrac calculation was created based on a distance matrix of all representative genomes compared to each other using Mash ^45^. Shannon diversity was calculated using skbio.diversity.alpha.shannon.

IgA+ probability thresholds for distinguishing “high binding”, “low binding”, and “uncoated” species were established based on manual inspection of the rank-sorted distribution of values (**Figure 2A**). To assess associations between particular taxa and IgA binding, at both the genus and phylum level, the scipy.stats and statsmodels python libraries were deployed. Only taxa with a minimum of 10 detections across all samples were considered in the analysis. For each taxa, we employed the Mann-Whitney U test to evaluate differences in IgA binding of each taxa versus all other taxa combined. The test was performed using the mannwhitneyu function from the scipy.stats package with a two-sided alternative hypothesis ^46^. To quantify the magnitude of the observed effect, the rank-biserial correlation was computed as the effect size for each Mann-Whitney U test. Fisher’s exact test was employed to investigate the association between each taxa and the three individual IgA binding classes. To calculate p-values and odds ratios, the 2x2 contingency table for each taxon and IgA binding class was constructed and passed to the fisher_exact function from the scipy.stats package. The false discovery rate (FDR) was controlled using the Benjamini-Hochberg procedure. The multipletests function from the statsmodels.stats.multitest package was applied to all p-values, originating both from the Mann-Whitney U tests and the Fisher’s exact tests, to obtain FDR-adjusted p-values.

To evaluate whether genera classified as “probionts” have higher IgA binding affinity (refer to the main text for the definition of “probionts”), we performed a focused analysis. For the purposes of this experiment, species were considered probionts if they belong to one four genera identified as candidates for next generation probiotics (*Bifidobacterium*, *Faecalibacterium*, *Akkermansia*, and *Eubacterium*) ^29,30^. We classified each species as probiont or not based on this definition, and we then computed the proportion of detections where the species exhibited either “high” or “low” IgA binding. This analysis was limited to species that were detected in at least 10 samples. We then used the rank-sum test from the scipy.stats package to compare these proportions between probiont and non-probiont species in aggregate (**Figure 2C**).

A phylogenetic tree was made from all species in the representative genome database (construction described above) using GToTree (v1.5.36) with default settings and the “Universal” gene set ^47^. The python program “treeswift” was then used to subset the tree to only include species with ≥ 5 detections ^48^. Heatmap visualization (**Figure 2D**) was performed using iTol ^49^.

### Gene annotation

We annotated all microbial genes detected in this study (n=1,171,838 robustly detected microbial genes) against four databases: Pfam (protein families) ^50^, KOfam (KEGG orthologs) ^51^, CARD (antibiotic resistance genes) ^52^, and the CAZy database (carbohydrate-active enzymes (CAZymes))^53^. Pfam annotations were performed using HMMer with the commands “hmmsearch --cut_ga -- domtblout --acc Pfam-A.hmm” and “cath-resolve-hits.ubuntu14.04” against the Pfam v32.0 database. KOfam annotations were performed using kofam_scan on default settings with the v103 database. CARD annotations were performed using the command “diamond blastp -f 6 -e 0.0001 -k 1” against the v3.2.5 database. CAZy annotations were performed using the command “hmmscan --domtblout Delta.faa_vs_dbCAN_v11.dm dbCAN-HMMdb-V11.txt GenomeSet_delta.genes.faa ; sh /hmmscan-parser.sh Delta.faa_vs_dbCAN_v11.dm > Delta.faa_vs_dbCAN_v11.dm.ps ; cat Delta.faa_vs_dbCAN_v11.dm.ps | awk ’$5<1e- 15&&$10>0.35’ > Delta.faa_vs_dbCAN_v11.dm.ps.stringent” with the dbCAN v11 database. More detailed instructions on gene annotation and parsing can be found here: https://instrain.readthedocs.io/en/latest/user_manual.html#gene-annotation. CAZyme substrate annotations were based on previously-used definitions ^54^.

### Machine learning

Gene annotation and abundance data was generated using the inStrian parse_annotations module (https://instrain.readthedocs.io/en/latest/user_manual.html#parse-annotations) with default settings. The resulting file “LongFormData.csv” was then parsed into a format acceptable for training with a machine learning classifier. Specifically, a table was created in which each row represented a genome detected in a sample, each column was an annotation, and values were “1” if the genome had the gene detected in either the IgA+ or native fraction of that sample, or “0” otherwise. A similar table was created based on the taxonomy of each organism, where the pandas method “pandas.get_dummies” was used to one-hot encode the taxonomy of each organism into a series of binary columns. This process effectively transformed the categorical taxonomy data into a format suitable for machine learning algorithms, where each unique category is represented by its own binary column. Finally, a merged table was created combining the columns of both tables into a single, merged table, based on each species detected in each sample.

The same machine learning training and evaluation procedure was used regardless of the input data source. First, class imbalance was handled by undersampling using the imblearn method “RandomUnderSampler”. Next, 20% of the data was removed for testing using the sklearn method “train_test_split”. A random forest classifier was then trained using the method “RandomForestClassifier” from sklearn, and overall accuracy was measured using the sklearn “accuracy_score” method. Receiver operating characteristic (ROC) curves were created using the sklean “roc_curve” method, area under the curve (auc) was calculated using the sklean “auc” method, and confusion matrices were created using the sklearn “confusion_matrix” method.

### Gene annotation statistics

Two statistical frameworks were deployed to associate specific microbial genes with IgA coating. The first framework uses Fisher’s exact test to identify annotations significantly associated with high IgA coating (**Figure 3D, E (left))**. The second framework uses Wilcoxon rank-sum tests to identify annotations associated with significantly higher IgA+ probability scores (**Figure 3G, E (right); Supplemental Figure S8)**. The two approaches yielded broadly consistent findings **(Supplemental Table S2)**.

#### Statistics based on Fisher’s exact test

The table “LongFormData.csv”, created as described above, was subset to only include genomes detected in either the IgA+ or native metagenomes. A contingency table was made for each annotation, representing: [[# of genomes with this annotation and “high” binding, # of genomes with this annotation without “high” binding],[# of genomes without this annotation and “high” binding, # of genomes without the annotation and without “high” binding]]. Annotation presence or absence for a genome in a fecal sample was based on the native metagenome. For higher taxonomic levels (e.g. genus and phylum) the same contingency tables were used, but subset to only include genomes from that particular taxa, and only taxa with at least 5 detections with “high” and at least 5 detections without “high” binding were considered. Fisher’s exact test was applied to each contingency table using “scipy.stats.fisher_exact”, and Benjamini-Hochberg FDR correction was performed using “statsmodels.stats.multitest”. The “effect size” or “IgA association” was calculated as the percentage of microbes with “high” IgA coating that encode this function - percentage of microbes with “high” IgA coating that do not encode this function.

#### Statistics based on Wilcoxon rank-sum test

The table “LongFormData.csv”, created as described above, was subset to only include genomes detected in either the IgA+ or native metagenomes. A Wilcoxon rank-sum test was performed for each annotation comparing 1) the IgA+ probability values of all detected genomes with this annotation, and 2) the IgA+ probability values of all detected genomes without this annotation. Annotation presence or absence for a genome in a fecal sample was based on the native metagenome. Only annotations detected at least 5 genomes, and not detected in at least 5 genomes, were considered. For higher taxonomic levels (e.g., genus and phylum) the same comparison was used, but subset to only include genomes from that particular taxa. Wilcoxon rank-sum tests were performed using the “scipy.stats.ranksums” command, and Benjamini-Hochberg FDR correction was performed using “statsmodels.stats.multitest”. The “effect size” or “IgA association” was calculated as the median IgA+ probability of organisms with the annotation - the median IgA+ probability of organisms without the annotation.

### *In situ* growth rate analysis

iRep values were calculated using the command “inStrain profile”, described above. iRep values were compared using the Wilcoxon signed-rank, as implemented through “scipy.stats.wilcoxon”. iRep values greater than 5 or less than 1 were considered invalid and excluded from this analysis. The paired test only considered genomes detected in both the native and IgA+ fractions, and among these genomes, the iRep values from the native samples were compared to those from the IgA+ samples.

### Association with multi-omic immune and health data

Previously generated multi-omic immune and health data ^25^ was downloaded and parsed into a single dataframe. The IgA binding, relative abundance in the IgA+ sample, and relative abundance in the native sample of each species in each sample was parsed into a single dataframe. Spearman’s rank correlation was then used to assess how strongly these microbial metrics (IgA binding, relative abundance in the IgA+ sample, and relative abundance in the native sample) are associated with the multi-omic immune, health, and microbiome data. The test was performed on the phylum level by including the data from all microbes of that phylum, and each individual immune / health data variable was tested independently. Tests were only performed when at least 10 samples exist where 1) at least one microbe from that phylum is detected in that sample and 2) that sample has the immune / health data being evaluated. Benjamini-Hochberg FDR correction was performed using “statsmodels.stats.multitest” within each of the 8 immune / health data categories. Heatmap visualization was generated using seaborn.

## Supplemental Note S1

An essential component of IgA MIG-Seq is processing the native and IgA+ fractions in identical manner. As detailed in the IgA MIG-Seq protocol, this includes processing the native fractions alongside IgA+ fractions until the cell-sorting step (including resuspending the native fecal samples in PBS), using the same DNA extraction and library preparation kits, and processing native and IgA+ fractions samples concurrently. This ensures that any differences between the sequencing data generated from these fractions is associated with IgA coating, and not due to differences in sample processing or batch effects.

To highlight this point, we compared the IgA+ fraction sequencing generated in this study to the sequencing of native fecal samples performed for previous publication ^25^ **(Supplemental Note S1 Figure 1**). These metagenomes were generated years prior, without resuspension in PBS, and using different DNA extraction and library preparation kits. The striking difference between those metagenomes and the native fractions sequenced in this study emphasize the need for sample processing uniformity. IgA MIG-Seq should not be done as an “add-on” to previously generated sequencing data.

**Supplemental Note S1 Figure 1:**
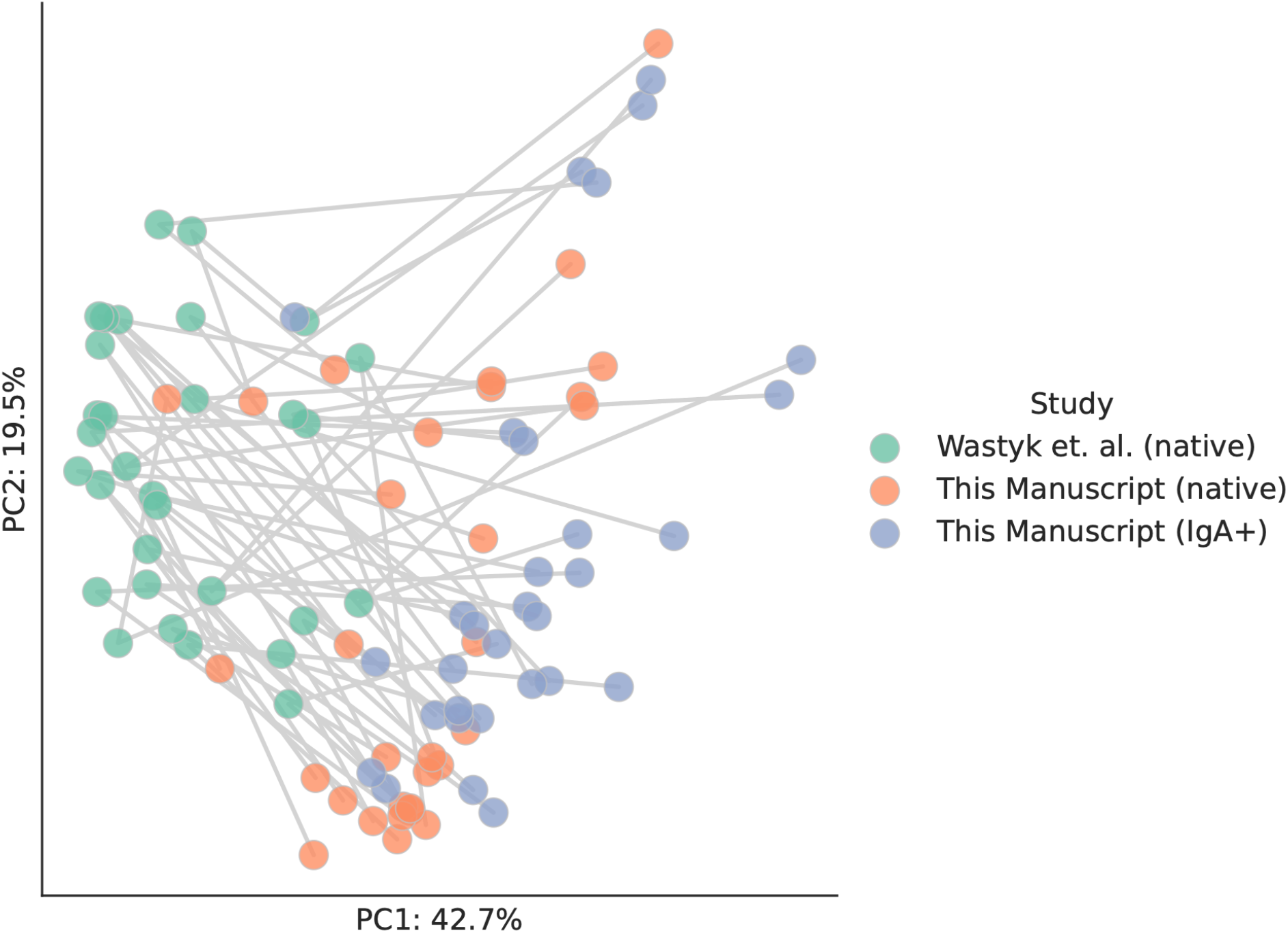
It is essential that native and IgA+ fractions are processed in identical manner. Each dot represents a metagenome from Wastyk et al. ^25^ (green) or this study (orange and blue). PCA is based on weighted UniFrac distance based on species-level relative abundance. All metagenomes were computationally processed in the same manner. Lines connect metagenomes generated from the same stool sample. Differences in sample processing including sequencing depth lead to large differences in reported microbiome composition (green vs. orange dots), highlighting the need for identical physical processing and sequencing of native and IgA+ fractions to identify accurate microbial IgA binding levels.

## Supplemental Note S2

Earlier studies have claimed to have performed metagenomic IgA sequencing, yet none have achieved the key milestones of the approach described in this study. Conrey et al. and other studies purport to employ “metagenomic sequencing”, but in reality the authors use the term “metagenomics” to refer to 16S sequencing ^4,55^. James et. al. performed shotgun metagenomic sequencing of IgA+ cell fractions, but the authors did not perform comparative sequencing on paired native fractions, substantially limiting the utility of the data ^56^. Fadlallah et al. performed shotgun metagenomic sequencing on both paired native and IgA+ fractions ^5^. However, the exceeding low sequencing depths and read lengths substantially limit data interpretation (approximately 0.6% of the sequencing depth applied in this study).

